# Publication games: in the web of reciprocity

**DOI:** 10.1101/2022.06.15.496272

**Authors:** Zoltán Barta

## Abstract

The present processes of research assessment, i.e. focusing on one or a few, related, scientometrics, foster questionable authorship practices, like gifting authorship to non-contributing people. An especially harmful one of these unethical practices is the formation of publication cartels, where authors offer gift authorship to each other reciprocally. Here, by developing a simple model and a simulation of the publication process I investigate how beneficial cartels can be and what measure can be used to restrict them. My results indicate that publication cartels can significantly boost members’ productivity even if paper counts are weighted by the inverse of author number (the 1/*n* rule). Nevertheless, applying the 1/*n* rule generates conflicts of interest both among cartel members themselves and between members and non-members which might lead to the self-purification of the academic publishing industry.

## 2 Introduction

Research integrity (ethical behaviour, sound methodology and rigorous peer review; Szomszor and Quaderi 2020) provides assurance that scientific activities lead to trustable and replicable results. Research integrity is, however, under threat as a result of how science currently operates. The recent, unprecedented expansion of science, exemplified, for instance, by the exponentially growing number of scientific articles (Fire and Guestrin 2019), gives way to the wide-spread use of scientometry for assessing the productivity and impact of researchers (Aubert Bonn and Bouter 2021). As science is usually funded by public resources, the desire to measure the performance of its actors is well justified. Introducing the assessment of scientists by one or a few metrics, like number of publications or citations, together with the hyper-competitiveness of science had, however, somehow unexpected consequences (Biagioli et al. 2019).

As, among others, Charles Goodhart observed, if a metric is used as a target then it becomes a bad metric (Edwards and Roy 2017; Fire and Guestrin 2019). This happens because people, in response to introduction of a target, alter their own behaviour to affect the metric directly instead to modify the activity the change of which was intended by introducing the metric (Werner 2015). In the recent process of corporatisation of science two such metrics became relevant: the numbers of papers and citations (Grossman and DeVries 2019).

Goodhart’s law is well illustrated by the introduction of the number of papers as a measure of productivity in science. Using this measure is based on the assumption that characteristics of scientific papers (like length or number of coauthors) are fixed and hence targeting more papers automatically leads to the generation of more new knowledge. Unfortunately, this was not what had happened, scientists responded in some unexpected, nevertheless clearly rational but sometimes unethical, ways (Fong and Wilhite 2017; Gopalakrishna et al. 2021). For instance, they reduced the length of papers (Fire and Guestrin 2019), i.e. they are publishing the same amount of knowledge in more papers (salami articles). Furthermore, mangling with authorship appeared where offering authorship to those who did not contributed to the given paper considerably (honorary authorship) can quickly increase their number of publications, again without any increase in knowledge produced (Aubert Bonn and Bouter 2021; Biagioli et al. 2019; Fong and Wilhite 2017; Gopalakrishna et al. 2021). A possible sign of this questionable authorship practice can be the recent raise of number of authors per paper (Fire and Guestrin 2019). One may argue that more authors per paper is the sign of science becoming more interdisciplinary. A recent analysis is, however, unlikely to support this conclusion; the number of coauthors increases with time even after controlling for attributes related to complexity of science (Papatheodorou, Trikalinos, and Ioannidis 2008). Another reason for the increased number of coauthors might be the increased efficiency that can follow from the increased possibility for division of labour facilitated by more authors (de Mesnard 2017). In this case, however, it is expected that the number of papers per author also increases, which seems not to be the case (Fire and Guestrin 2019).

Questionable authorship practice, on the other hand, appears to be common. Recent surveys suggest that about 30% of authors were involved in these unethical practices (Biagioli et al. 2019; Fong and Wilhite 2017; Gopalakrishna et al. 2021; Grossman and DeVries 2019; Halaweh 2020; Marušić, Bošnjak, and Jerončić 2011). One of these practices is ghost authorship when someone who has significantly contributed to the article is excluded from the author bylist (Jabbehdari and Walsh 2017). In other forms (honorary authorship) just the opposite happens; those are offered authorship who have not (considerably) contributed to the work published (Fong and Wilhite 2017; Gopalakrishna et al. 2021). Several reasons can be behind gifting authorship to someone. Junior authors might include more senior ones because of respect or they are forced to do so (Pan and Chou 2020). Senior authors may gift authorship to juniors in order to help them obtain post-doctoral scholarships or tenure (Von Bergen and Bressler 2017).

A very efficient way to increase the number of publications may be to practice honorary authorship reciprocally. The most organised form of this behaviour is founding publication cartels. The cartel is formed by a group of people who agree to mutually invite each others to their own publications as guest authors without any contribution. As in recent assessment practice coauthored papers count as a whole publication to every coauthor on the bylist, publication cartels can significantly boost the productivity of cartel members. This is the phenomenon which is called as ‘publication club’ by de Mesnard (2017). As the noun of ‘club’ involves a positive connotation I prefer to use ‘cartel’ for this under studied but highly unethical behaviour. Simple argument suggests that sharing the credit of a publication among the coauthors can decrease the incentive of forming cartels (de Mesnard 2017). The simplest scenario for sharing is the 1/*n* rule under which only 1/*n* part of a publication is attributed to each of the *n* coauthors of the given paper (de Mesnard 2017).

In this paper I develop a simple model of publication cartels to understand how effective they are to increase members’ productivity and whether it is possible to eliminate them by applying different measures, like the 1/*n* rule. I then extend my study to situations resembling more to real world conditions by developing a computer simulation of cartels. I use this simulation to investigate how using different metrics of productivity affect authors outside of cartels.

## 3 The model

We compare the publication performance of two authors, author *A*_1_ and author *B*_1_. Authors work in separate groups (group *A* and group *B,* respectively) each of which contains *G*_i_ (*i* = *A* or *B*) people (including the focal author). Each author in group *A* produces *p*_A_ papers in a year by collaborating with *c*_A_ authors from outside of the group, i.e. their primary production is *p*_A_. Similarly, each author in group *B* primarily produces *p*_B_ papers by collaborating with *c*_B_ people outside of the group. The difference between authors *A*_1_ and *B*_1_ is that authors in group *A* work independently of each other, while authors in group *B* invite all other group members to be a coauthor on their papers independently of their contribution to that paper (Fig 1). In other words, authors in group *B* form a publication cartel.

**Figure 1:**
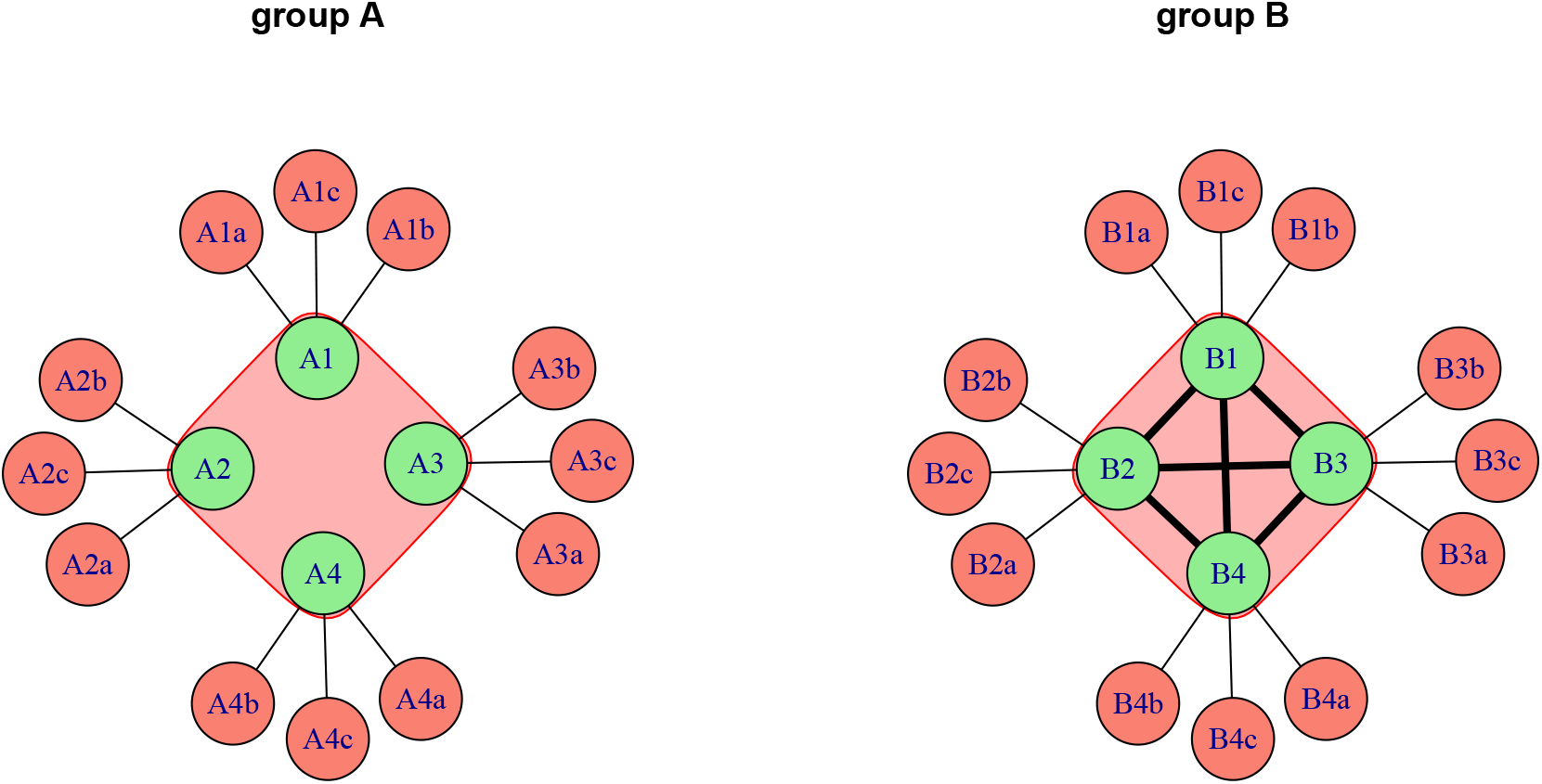
The publication relationships in groups *A* and *B* of the model. Nodes are authors, while edges symbolise shared publications. Groups of four authors are marked by the underlying shapes. In group *A* authors work with several coauthors from outside of the group but they do not invite group mates to be coauthors on their own papers. Contrarary, authors in group *B* form a publication cartel i.e. each author invites all other authors in the group to be a coauthor (note the connections between group members).

For simplicity, we assume that *G*_A_ = *G*_B_ = *G* (*G* > 1), *p*_A_ = *p*_B_ = *p* and *c*_A_ = *c*_B_ = *c*, i.e. author groups are of the same size, authors produce the same number of primary papers and they have the same number of coauthors from outside of the group. In this case the total numbers of papers produced by the groups, the group productivity, are equal (*Gp* = *G*_A_*p*_A_ and *G*_B_*p*_B_, respectively). The total numbers of papers (co)authored by authors *A*_1_ and *B*_1_ are, however, different. Author *A*_1_ writes *n*_A_ = *p*_A_ = *p* papers. On the other hand, author *B*_1_ (co)authors *n*_B_ = *p*_B_ + (*G*_B_ - 1) *p*_B_ = *G*_B_ *p*_B_ = *Gp* papers. In the case of author *B*_1_ the term (*G*_B_ - 1)*p*_B_ represents the papers on which author *B*_1_ is invited as honorary author. It is easy to see that as far as *G* > 1, author *B*_1_ will have many more paper than author *A*_1_, i.e *n*_B_ > *n*_A_.

A natural way to correct for this bias is to taking into account the number of authors each paper has and instead of counting the papers themselves as a measure of productivity one sums the inverse of the number of authors (the 1/*n* rule, de Mesnard 2017; Vavryčuk 2018):

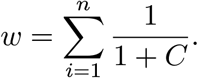

Here, number 1 in the denominator symbolises the focal author, while *C* is the number of coauthors. For author *A*_1_, *C* = *c*_A_ = *c*. On the other hand, for author *B*_1_, *C* = (*G*_B_ - 1) + *c*_B_ = (*G* - 1) + *c*. If *c* = 0, then the division by the number of coauthors works, we regain the number of papers the authors produced without inviting their group members.

For author *A*_1_:

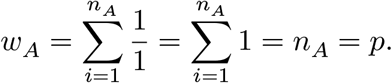

For author *B*_1_:

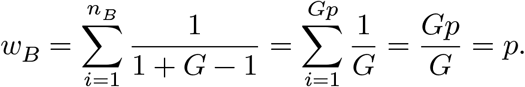

On the other hand, if the focal authors collaborate with others outside of their groups, as Fig 1 illustrates, the situation changes (Fig 2):

For author *A*_1_:

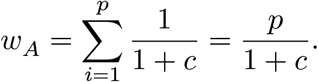

For author *B*_1_:

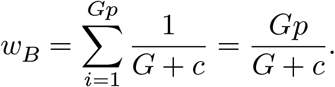

The weighted number of papers produced by author *B*_1_ relative to author *A*_1_, *w*_B_/*w*_A_, is:

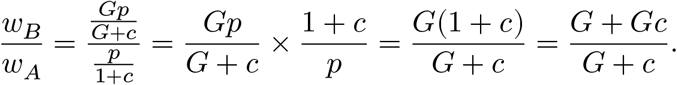

The proportion of *w*_B_/*w*_A_ is greater than one if *G*+*Gc* > *G*+*c*, which is always true if *c* > 0 (as we already assumed *G* > 1; Fig 2). This means that if authors collaborate anyone from outside of their groups then authors in group *B* will always have higher publication performance than authors in group *A*, despite the fact that the two groups have the same productivity.

**Figure 2:**
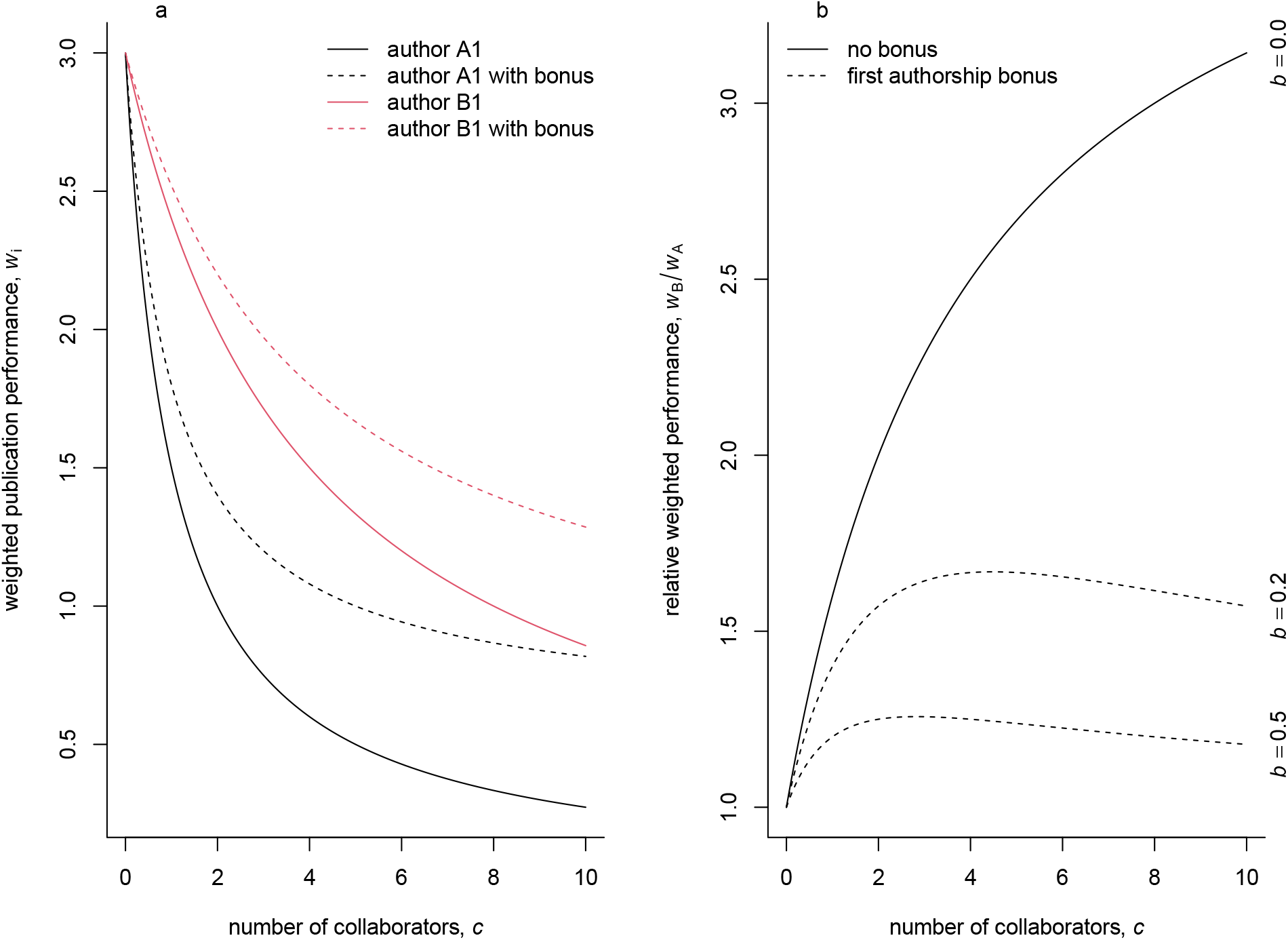
Publication performance when authors collaborate with people from outside of their groups. Weighted publication performance of authors *A*_1_ and *B*_1_ (a). Weighted publication performance of author *B*_1_ relative to that of author *A*_1_ (b). The weighted publication performance is calculated by taking into account the number of coauthors. During this calculation first authorship can be rewarded by a bonus, *b.* If *b* = 0, then each coauthors receive the same weight for a given publication. On the other hand, if *b* >0, the weight of the first author is higher then that of the coauthors, i.e. the first author of a paper is rewarded. On subpanel (a) *b* = 0.2, on (b) *b* is given on the right margin.

To compensate for this productivity bias, author *A*_1_ should produce *w*_B_/*w*_A_ times more papers, *p*_A_ = *p*_B_(*G* + *Gc*)/(*G* + *c*). This surplus of papers needed for compensating the productivity bias increases with *c* and it keeps to *G*.

Authors in group *A* can also compensate for the productivity bias by decreasing the number of their collaborators from outside of the group. This reduction must be by a factor of *G*: *c*_A_ = *c*_B_/*G*.

A useful modification to the 1/*n* rule is the so called *first-author-emphasis* scheme (Vavryčuk 2018). In this scheme, the first authors receive a bonus, *b,* to recognise their leading role in producing the papers. Under this scheme the weighted publication performance for author *A*_1_, *w*_A_’, is:

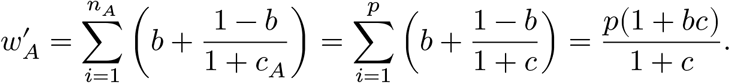

Here, the first author, who is author *A* for all his papers, get a bonus *b* for contributing most to the paper, and the rest of the credit, 1-*b*, is divided equally between all authors (including the first author, Vavryčuk 2018). The weighted publication performance for author *B*_1_ under the first author scheme, *w*_B_’, is:

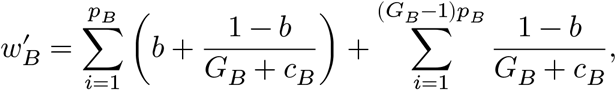

where the first term gives the credit for first author papers, while the second one is for the coauthored papers. After simplification, we obtain:

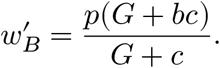

By comparing *w*_B_’ to *w*_A_’ it is easy to show that author *B*_1_ will always have a higher publication performance than author *A*_1_, i.e. *w*_B_‘/*w*_A_’ > 1, if *G* >1 and *b* <1. Further analysis,

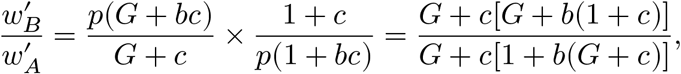

shows that for *w*_B_‘/*w*_A_’ > 1, the condition *c* > 0 should also be fulfilled. As numerical computation indicates (Fig 2) the bias is decreased by introducing the first authorship bonus, but it is still significant. Vavryčuk (2018), for instance, recommend a bonus of *b* = 0.2, but in this case author *B*_1_ sill has around 50% more credit for the same work than author *A*_1_ has. The difference between authors *A*_1_ and *B*_1_ decreases as *b* increases (Fig 2b), but this way coauthorship is worth less and less, undermining the possible benefits of collaborations.

To summarise, this simple model shows that the formation of publication cartels can be an advantageous, but unethical, strategy to increase publication productivity even if one control for the number of coauthors of papers. Note, however, that this model might be overly simplified as all authors have the same primary productivity and we do not investigated how productivity of authors outside of the cartels changes as a consequence of founding cartels. To obtain a more realistic understanding of publication cartels next I develop a simulation of the publication process.

## 4 The simulation

We start simulating the publication process with constructing a publication matrix of papers and authors, *M*_P_ (Fig 3). Element *a*_ij_ of *M*_P_ is one if author *j* is on the bylist of paper *i* and zero otherwise. Therefore, *M*_P_ can be considered as a matrix representation of a bipartite graph, where rows and columns represent the two types of nodes, papers and authors, respectively. To construct *M*_P_ we consider a community of *c* authors. The number of papers written by author *j* in the community is given by *k*_j_. For the community we construct an empty matrix (all *a*_ij_ = 0) of size *p* and *c*, where *p* > max(*k*_j_). Then for each column *j* we randomly distributed *k*_j_ number of ones over the *p* empty places. Having constructed *M*_P_ we create a weighted collaboration (or co-authorship) matrix, *M*_C_, by projecting *M*_P_ to the nodes of authors. The weights of *M*_C_, *J*_ij_, are Jaccard similarity indices calculated between each pair of authors *i* and *j* (*i* ≠ *j*) as

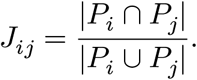

**Figure 3:**
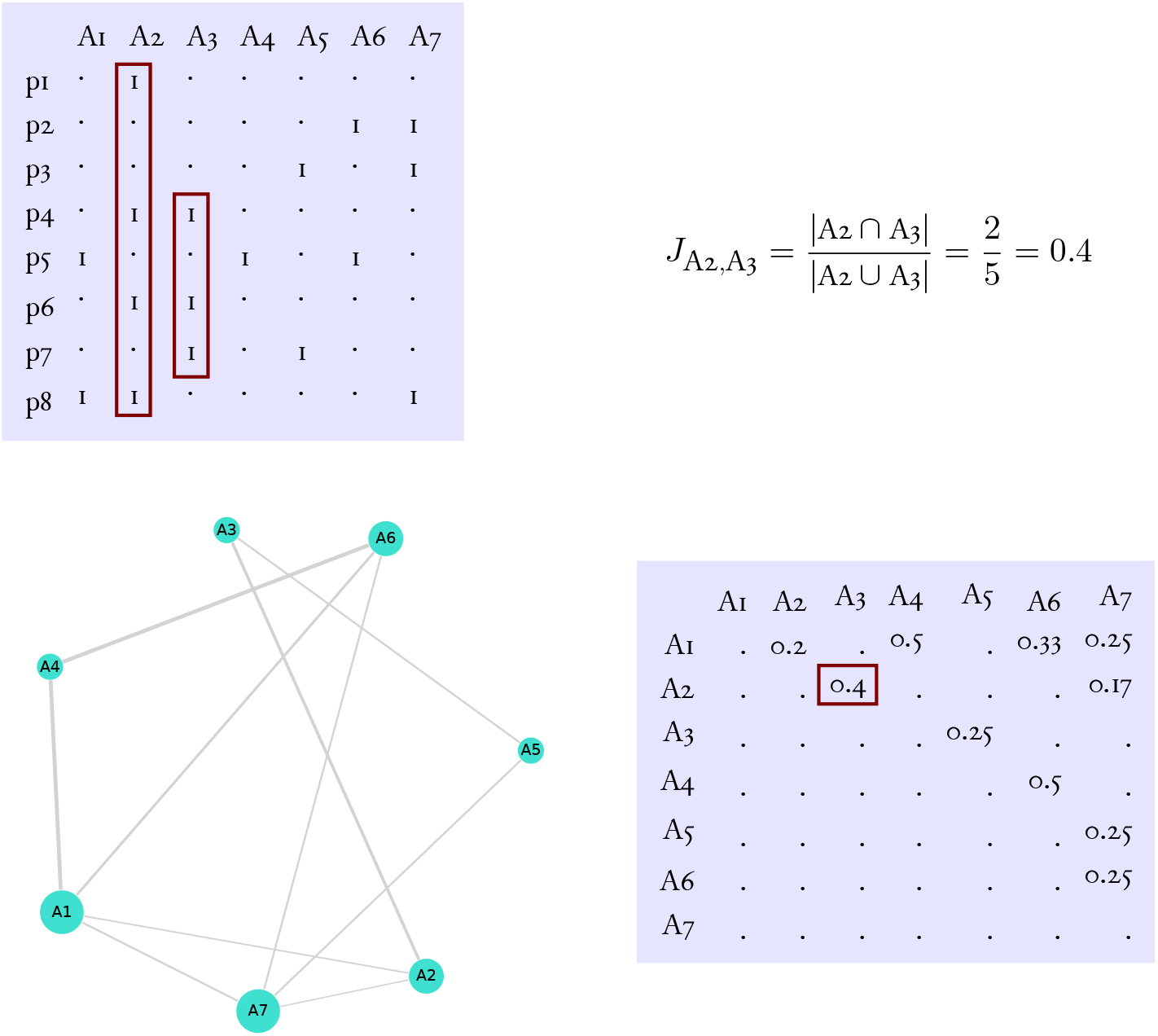
The construction of publication network. The top left panel shows the publication matrix, *M*_P_. Each row and column of this matrix represents a paper and an author, respectively. Values of 1 indicate that an author is on the author list of a given paper, while dots symbolise zeros. From the publication matrix one can derive the collaboration matrix, *M*_C_ (bottom right panel) by calculating the Jaccard simmilarity (top right) for each possible pairs of authors. The bottom left panel shows the resulting weighted, undirected collaboration graph, *G*_C_. The red rectangles exemplifies the calculation of Jaccar simmilarity.

Here, *P*_i_ is the set of papers to which author *i* contributed. In other words, the weight between two authors is the proportion of shared papers to the total number of unique papers to which either of authors *i* or *j* contributed to. It varies between zero (i.e. no common publication between author *i* and *j*) and one (i.e. all publications by the two authors are shared). Note, Jaccard similarity between authors in group A of the above model is zero, while between authors in group B is one. From *M*_C_ we construct a collaboration graph, *G*_C_.

I simulated the formation of cartels by choosing |*κ*| authors from the community (Fig 4). Let *κ* is the set of cartel members. Then, with probability *p*_c_, I changed each element *a*_ij_ = 0 of *M*_P_ to *a*_ij_ = 1 where the following conditions met: *j* ∈ *κ* and at least one *a*_ik_ = 1 with *k* ∈ *κ* but *k* ≠ *j*. I project the resulting publication matrix, *M*_P_’ to *M*_C_’ and constructed the corresponding collaboration graph, *G*_C_’.

**Figure 4:**
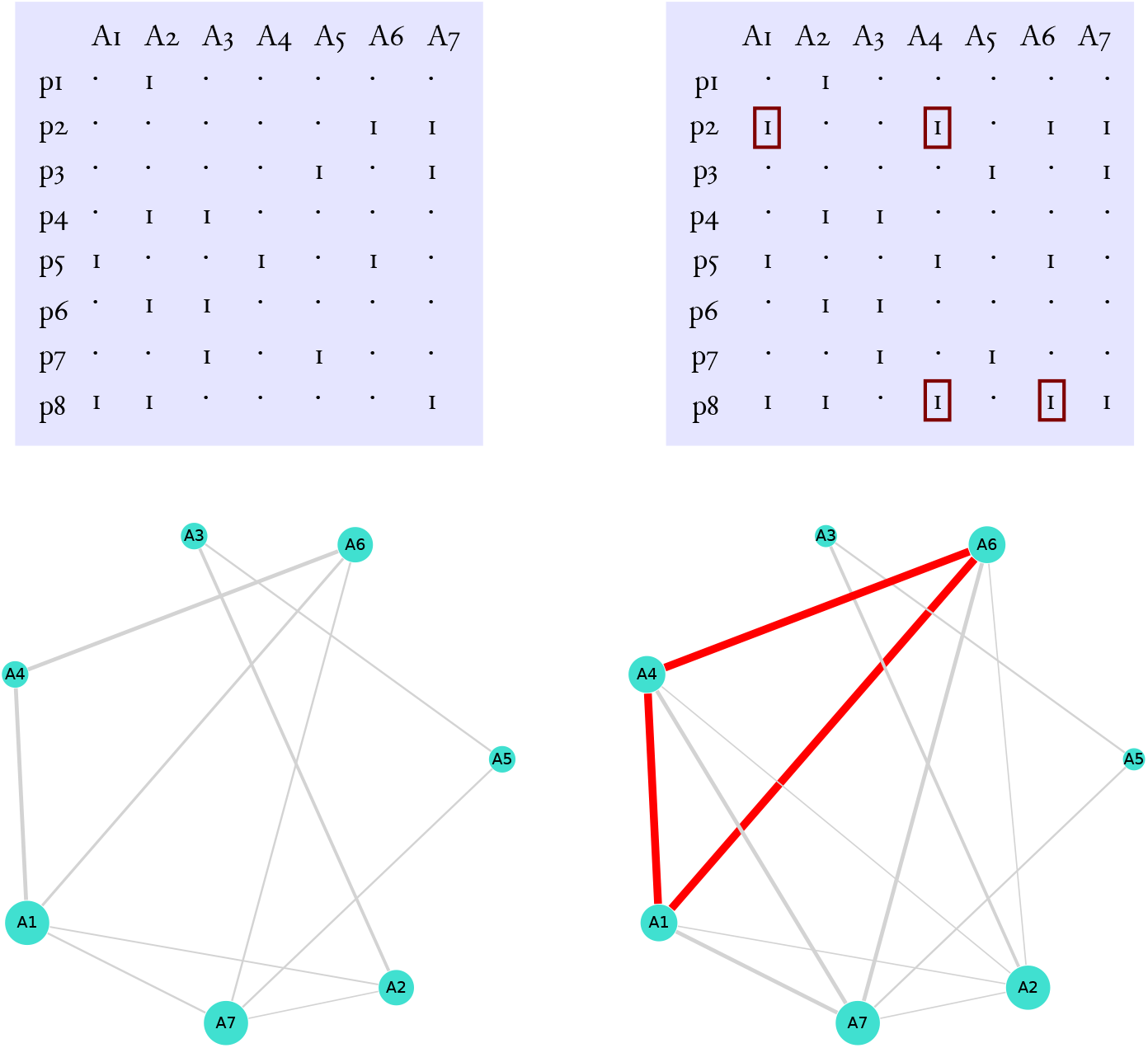
The formation of cartels. The panels on the left illustrate a publication network without cartel. The panels on the right show how a cartel between authors A1, A4 and A6 can be formed: Author A6 invites authors A1 and A4 to be coauthors on paper p2, while author A1 do the same with authors A4 and A6 on paper p8. The small red rectangles mark the authorships gained this way. The bottom right panel shows the resulting collaboration graph, where the red edges connect cartel members. Note (i) the strong connections between members and (ii) adding cartels also changes the connections of non-members.

By setting all *k*_j_ = *k* and *p* > > *k* we can simulate the case of equal productivity and no collaboration from outside of the group. Here the simulation produces the same results as the model: productivity of cartel members increased but this can be accounted for by using weighted number of publications.

To induce collaboration between authors I next set *p* < ∑*k* (the authors still have the same productivity prior to cartel formation). Under these conditions, if we consider the number of papers, the productivity of cartel members increases significantly by forming cartel while productivity of non-members does not change (Fig 5). In accordance with the model, the productivity of cartel members increases even if we consider the weighted number of papers. Interestingly, the productivity of many non-members decreases when cartel is formed (Fig 5).

**Figure 5:**
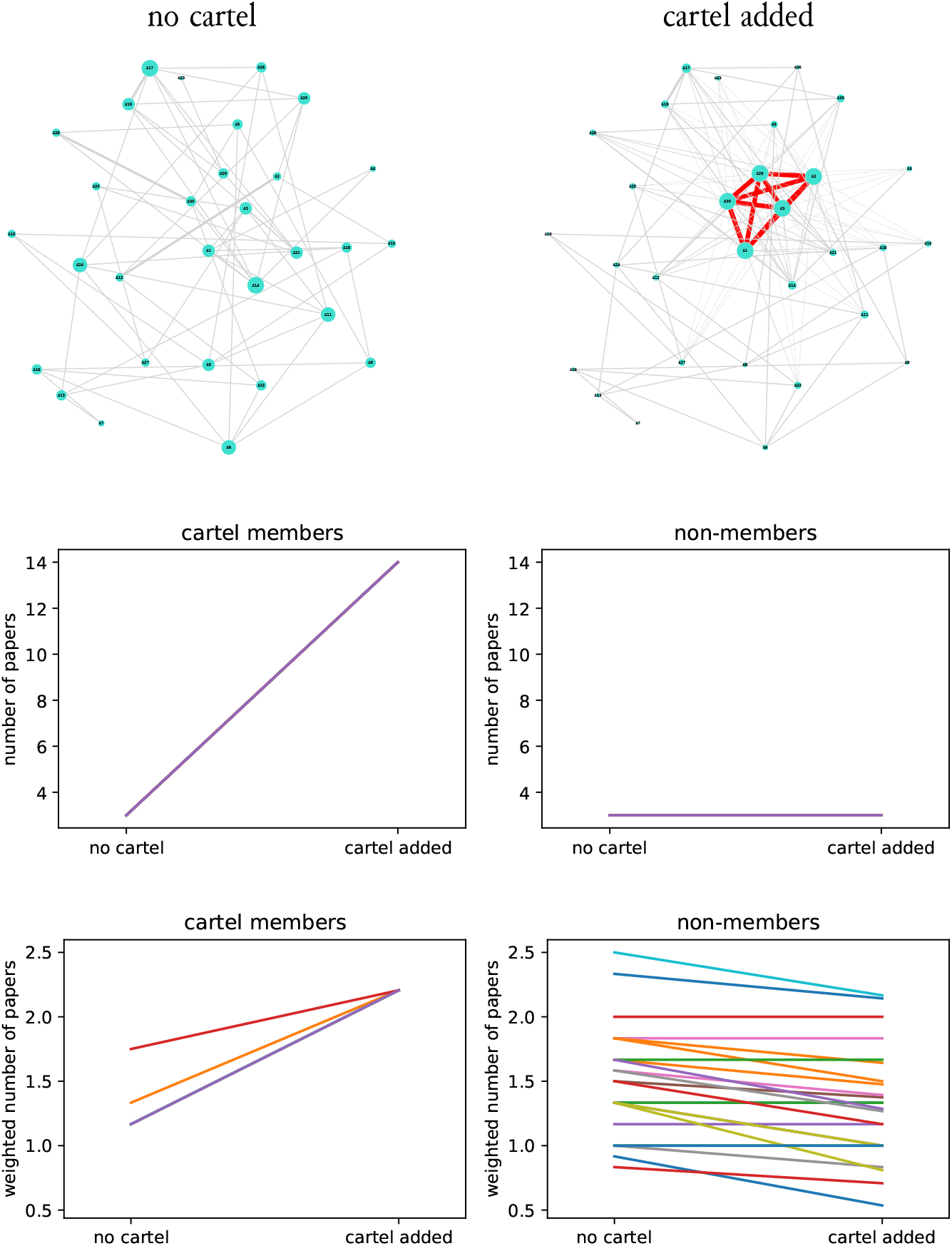
The effect of cartel formation on the productivity of cartel members and non-members: equal prior productivity of authors. The top panels illustrate the collaboration graph before and after cartel formation. The middle panels show how the number of papers produced by members and non-members changes because of founding cartel. The bottom panels illustrate the same but using the weighted number of papers as a measure of productivity. Collaboration graph formed with *c* = 30, *k* = 3, *p* = 60, *p*_c_ = 1 and *κ* = {1, 2, 3, 29, 30}.

I further generalise the simulation results by setting the prior productivity of authors to different values (Fig 6). Using the number of papers as metric leads to the same conclusions: members’ productivity increases after cartel formation, non-members’ productivity does not change. On the other hand, using the weighted number of papers reveal an interesting effect: the productivity of cartel members with high prior productivity have their productivity being decreased because of cartel foundation (Fig 6). Similarly to the previous case, productivity of many non-members decreases as a consequence of cartel formation (Fig 6).

**Figure 6:**
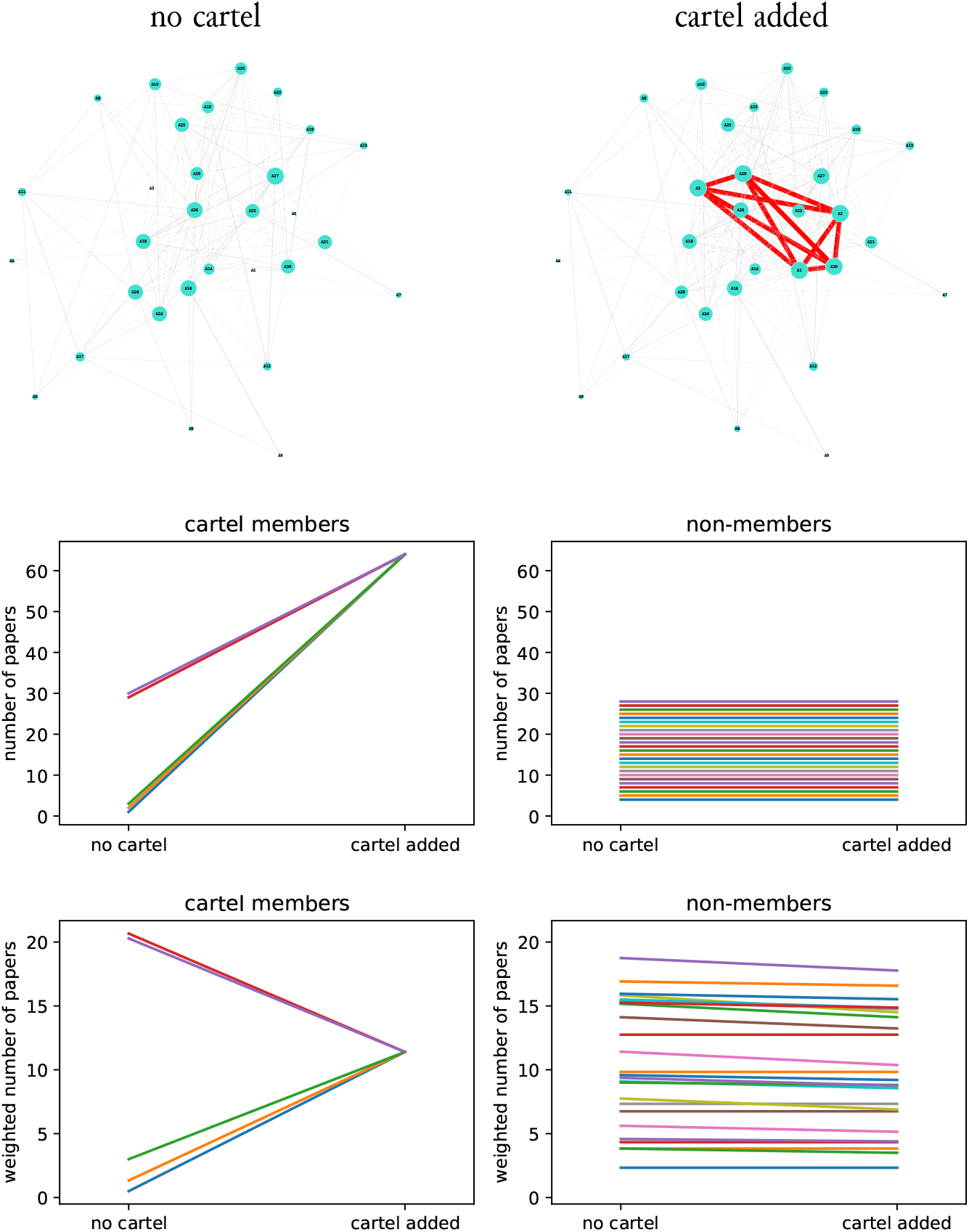
The effect of cartel formation on the productivity of cartel members and non-members: prior productivity of authors differs. The top panels illustrate the collaboration graph before and after cartel formation. The middle panels show how the number of papers produced by members and non-members changes because of founding cartel. The bottom panels illustrate the same but using the weighted number of papers as a measure of productivity. Collaboration graph formed with *c* = 30, *k*_jj_ = j, *p* = 60, *p*_c_ = 1 and *κ* = {1, 2, 3, 29, 30}.

## 5 Conclusions

Under the current climate of wide spread use of scientometry indices to assess academics publication cartels can provide huge, although unethical benefits. As my results indicate, members of cartels by reciprocally inviting each other as honorary authors can easily boost their own publication productivity, i.e. the number of papers they appear on as (co)author. As many scientometrics currently in use are strongly associated with the number of publications a scholar has produced (Aubert Bonn and Bouter 2021; Fong and Wilhite 2017; Grossman and DeVries 2019) becoming cartel member can have a very general positive effect on one’s academic career.

One may consider that fighting off cartels is not necessary because of “no harm no foul”: research integrity may not be inevitably damaged by cartel foundation, cartels can produce high quality research. Nevertheless, cartels do distort the research competition landscape. This might result in that highly competent, talented researchers, who are not members of any cartels, are forced into inferior roles which, in turn, compromises the society’s ability to produce more novel and innovative results. Therefore, cartel formation should be restricted.

Fighting against cartels is, however, not trivial. First, identifying cartels, not to mention to prove for a group of researchers that they are cartelling is inherently difficult. Investigating properties of coauthor networks might help. Nevertheless, a possible way to restrict cartels without their identification is to use such scientometrics which penalise cartel formation. An obvious choice can be to weight the number of publications an author has by the inverse of the number of authors on the bylists of these papers, the so-called 1/*n* rule. As my calculation shows this rule can only be effective if coauthorship only occurs between cartel members. As soon as collaboration is wide spread among both cartelling and non-cartelling authors my results indicate that the 1/*n* rule breaks down and cartel members still gain undeserved benefits. On the other hand, my computations also show that the 1/*n* rule can still be useful against publication cartels, because it generates conflicts of interest among the parties. Collaborators of cartel members suffer a loss if the 1/*n* rule is applied which might force them either to change the unethical behaviour of cartel members or abandon to collaborate with them.

The results of the simulation also suggest that, given that the 1/*n* rule is used to rate authors, cartel members should be of similar prior productivity, because cartel establishment with lowly ranked authors can have a detrimental effect on the productivity of prolific authors. This would also introduce a conflict of interest between authors leading to that founding cartels is only worth for low productivity authors among themselves, because (i) highly ranked authors do not need to manipulate their productivity indices (they are already high) and (ii) they can actually loose on cartel formation, given that productivity is measured by weighted number of papers, i.e. according to the 1/*n* rule.

To summarise, I strongly argue for using the 1/*n* rule as the basis of scientometry. Unfortunately, its general use is opposed by many parties for many reasons. It still remains to see whether these reasons are valid or not, but my calculations indicate that the application of 1/*n* rule can generate such processes which may ultimately lead to the self-purification of the academic publication industry. Of course, abandoning the current, metric-only research assessment system can also help.

## 6 Acknowledgments

I thank Miklós Bán, Gábor Lövei, Tibor Magura and Jácint Tökölyi to review a previous version of the manuscript. The research was supported by the Thematic Excellence Programme (TKP2020-IKA-04) of the Ministry for Innovation and Technology in Hungary.

## Notes

### Competing Interest Statement

The authors have declared no competing interest.

